# Hyperosmolar stress promotes the release of small extracellular vesicles containing metabolic proteins from corneal epithelial cells

**DOI:** 10.64898/2026.03.27.714594

**Authors:** Belinda J. Hernandez, Valentina Morakis, Andrew Lemoff, Anupam Mondal, Danielle M. Robertson

**Author notes:** Corresponding author: Danielle M. Robertson, OD, PhD, The Department of Ophthalmology, UT Southwestern Medical Center, 5323 Harry Hines Blvd, Dallas, TX, USA 75390-9057. Support: NIH/NEI grants EY024546 (DMR), Core Grant for Vision Research EY030413, a Challenge Grant from Research to Prevent Blindness, and the Shirley G. and Norman Alweis Endowment Fund for Vision (DMR).

## Abstract

**Purpose:** Hyperosmolar stress (HOS) is a major contributor to corneal epithelial cell damage in dry eye disease. We have previously shown that HOS damages mitochondria and impairs cell metabolism in corneal epithelial cells. Small extracellular vesicles (sEVs) are cell-derived lipid envelopes that are present in all body fluids, including tears. Prior studies suggest that sEV release and composition may be linked with changes in cell metabolism. In this study, we tested the effects of HOS on sEV release and composition, and found that sEV cargo may reflect early, underlying changes in dry eye disease.

**Methods:** Telomerase-immortalized human corneal epithelial (hTCEpi) cells were treated with 450 mOsm NaCl for five days to induce chronic HOS. sEVs were isolated using differential centrifugation followed by iodixanol density gradient flotation. Particle number was determined using Nanoparticle Tracking Analysis (NTA). Mass spectrometry was used to assess the sEV proteome, and selected proteins were validated by immunoblot. Proteome pathways were analyzed using KEGG and CORUM.

**Results:** Pathway analysis revealed an increase in metabolic proteins and proteasome components in sEV cargo released from hTCEpi cells exposed to HOS. These proteins were increased more than fourfold in HOS-sEVs. Examination of proteins involved in the endosomal pathway and NTA further confirmed an increase in HOS-sEV release.

**Conclusion:** Our findings suggest a potential mechanism whereby corneal epithelial cells exposed to HOS retain proteins involved in maintaining tissue integrity, while simultaneously releasing unneeded proteins involved in cell metabolism. The presence of metabolic proteins in sEVs may serve as early indicators of dry eye disease.

## Introduction

The cornea is a transparent avascular tissue that forms the front part of the eye. In addition to the refraction of light, the cornea serves as a first line of defense to protect the eye from opportunistic pathogens^1^. Due to its external location, the cornea is continuously exposed to stress from the outer environment, leading to corneal epithelial cell surface damage. In dry eye disease (DED), the disruption of the precorneal tear film, along with damage to surface corneal epithelial cells, can greatly impair vision. Hyperosmolar stress (HOS) is a major stimulus for corneal epithelial cell damage in DED, leading to inflammation, cell shrinkage, and perturbation of cell function^2, 3^. We have previously shown that HOS also impairs corneal epithelial cell metabolism^4^. In various cell systems, changes in cell metabolism have been associated with changes in extracellular vesicle (EV) formation, release, and composition.^5^

Extracellular vesicles are nano-sized vesicles enclosed within a lipid bilayer that are secreted by all cells in the body and play a key role in cell-to-cell communication^6^. There are three established subtypes of EVs: apoptotic bodies, microvesicles, and small EVs (sEVs)^6^. sEVs are cell-derived nanovesicles (30-150 nm in diameter), which are formed and released by the endosomal pathway^7^. The cargo of sEVs consists of proteins and genetic material (miRNAs/DNA) from their cell of origin. This enables cells to communicate in a non-contact manner and modulate immune-regulatory pathways and other critical cell processes^6, 8–10^. sEVs are released from most cell types during homeostasis and in disease states, consequently influencing the activity of recipient cells through the delivery of bioactive signals^6, 11^. Multiple lines of evidence have shown that corneal epithelial cells release sEVs and that EVs are present in the precorneal tear film^12–18^. Thus, EVs released by stressed or damaged corneal epithelial cells may serve as a fertile source of early indicators of DED.

In this study, we interrogated the role of sEVs released by corneal epithelial cells during isosmolar culture and HOS, the latter a known cause of cellular damage in the corneal epithelium. To accomplish this, we established a chronic subtoxic model to analyze the number and cargo of sEVs released from corneal epithelial cells before the onset of overt morphological changes. Importantly, we show for the first time that chronic subtoxic stress in corneal epithelial cells induces changes in the proteome cargo of sEVs. We further identified distinct changes in the metabolic proteins contained within sEVs released in response to HOS. Thus, the increase in metabolic proteins may serve as novel early indicators of corneal epithelial cell health.

## Material and Methods

### Cell Culture

A human telomerized corneal epithelial (hTCEpi) cell line previously developed and characterized by our laboratory was used for these studies^19^. hTCEpi cells were cultured in serum-free keratinocyte basal media (KBM) containing supplements (KGM, PromoCell, Fisher Scientific, Waltham, MA) and 10% penicillin-streptomycin-amphotericin B (Lonza, Walkersville, MD). Cells were maintained at a final CaCl_2_ concentration of 0.15 mM by supplementing KGM with additional CaCl_2_ (calcium chloride solution, 0.5 M, PromoCell). For sEV collection, hTCEpi cells were grown on T-175 flasks. After cells reached 100% confluency, media was changed to KBM (devoid of KGM supplements) containing 10% penicillin-streptomycin-amphotericin B (Lonza, Walkersville, MD), and 0.15 mM of CaCl_2_. Hyperosmolar stress (HOS) was induced by adding media containing 450 mOsm of NaCl to fully confluent hTCEpi cells. Cells were treated for five days. hTCEpi cells in isosmolar (330 mOsM NaCl) media were cultured in parallel. All cells were maintained at 37 °C and 5% CO_2_.

### sEV Isolation

For each sEV preparation, 260 to 270 mL of conditioned media were used as starting material. Media was collected daily and stored at −80°C until use. Samples were then thawed, and the conditioned media that were collected over the five days were pooled for test and control cells. Small EVs were isolated according to the MISEV2023 guidelines^20^ using cushioned iodixanol buoyant density gradient ultracentrifugation^21, 22^. Briefly, conditioned media samples were centrifuged at 2000 *g* for 10 minutes to remove cell debris. The resulting supernatant was collected and kept at −80°C. Upon thawing, cleared supernatants were concentrated using Corning Spin-X 20 mL centrifugal filter devices with a 100,000 molecular weight cutoff (Corning, Glendale, AR). The concentrated media samples were then placed in polyallomer tubes (Beckman Coulter Inc., Indianapolis, IN) and centrifuged at 10,000 g for 30 minutes using the Beckman XPN-100 Ultracentrifuge. The resulting supernatant was carefully placed in an Ultraclear X tube (Beckman Coulter Inc., Indianapolis, IN) onto a 2 mL cushion of 60% OptiPrep (Sigma-Aldrich, Saint Louis, MO) and centrifuged at 100,00 *g_avg_* in an SW32 Ti rotor for 180 minutes. The resulting 2 mL cushion and 1 mL of interface were collected from the bottom using a 4-inch blunt 18-gauge metal hub Luer lock needle (Hamilton Company), with a 5 mL syringe (Becton Dickinson, Franklin Lakes, NJ) extending from the top. The 3 mL containing the cushion and the gradient was placed at the bottom of a gradient tube (Beckman Coulter, Indianapolis, IN) and then used as a 40% OptiPrep fraction in the subsequent density gradient. A discontinuous gradient of iodixanol solution was then prepared by carefully overlaying the bottom fraction with 3 mL of 20%, 10%, and 5% solutions of iodixanol buffered with [0.25M sucrose, 10 mM Tris-HCl (pH 8.0)], respectively. The gradient tubes were centrifuged at 200,000 *g_avg_* in a SW 41 Ti Rotor for 4 hours at 8°C. One mL fractions were collected from top to bottom and weighed to determine density. The one mL fractions were then diluted with phosphate-buffered saline (PBS) and subjected to centrifugation at 100,000 *g_avg_* for 90 minutes in a SW 41 Ti rotor. Pellets were resuspended in 70-100 µL of RIPA buffer containing Protein Protease Inhibitor or in 200 µL PBS. Samples were stored in −80 °C for further analysis.

#### sEV Characterization

Total protein content in sEV preparations was determined using a BCA Pierce Protein Assay (ThermoFisher Scientific, Waltham, MA). Western blotting to test for proteins in sEVs and hTCEpi cell lysates was performed as previously described^23^. Briefly, sEV samples and whole cell lysates were electrophoresed on a 4-12% gradient gel (Bio-Rad, Hercules, CA). The proteins were blotted onto polyvinylidene difluoride (PVDF) membranes (Bio-Rad, Hercules, CA) using a BioRad Trans-Blot Turbo Semi-Dry transfer apparatus, and then probed with the indicated antibodies. Primary antibodies were incubated overnight at 4 °C. The antibodies used in this study include: Anti-CD9 (Abcam, Waltham, MA), Anti-gamma-Catenin (Junctional Plakoglobin, Cell Signaling, Danvers, MA), Anti-Desmoglein (BD Transduction, Franklin Lakes, NJ), Anti-Desmoplakin (Proteintech, Rosemont, IL), and Anti-MDH2 (Abcam). These antibodies were used at a concentration of 1:500. Anti-Annexin 2 (Abcam), Anti-Alix (Abcam), Anti-PGK1 (Abcam), Anti-LDH2 (Cell Signaling), Anti-Calreticulin (Cell Signaling), and anti-SOD1 (Proteintech), β-actin rabbit HRP conjugated (Cell Signaling), and α-tubulin (Cell Signaling) were used at 1:1000. Secondary antibodies, anti-Rabbit horseradish peroxidase (HRP) and anti-Mouse HRP were purchased from Bio-Rad (Hercules, CA). To assess particle concentration and size, a nanoparticle tracking analyzer (NTA) (NanoSight NS300, Malvern Panalytical, Worcestershire, UK) configured with a 532 nm laser and a high-sensitivity scientific CMOS camera was used. A total of five readings were obtained per condition and averaged to determine particle concentration analysis.

### Mass Spectrometry and Proteomics

An untargeted proteomics analysis was performed using three sets of sEV preparations from hTCEpi cells grown in media containing 330 mOsm and 450 mOsm. culture. Equal total protein concentrations for all samples were mixed with Laemmli buffer (Bio-Rad, Hercules, CA) and subjected to SDS-PAGE^24^. Samples were run approximately 1 cm into the separating gel^23^. Samples were digested overnight with trypsin (Pierce Biotechnology, Rockford, IL) at 37°C following reduction with 20 mM dithiothreitol (DTT) at 37°C for 1 hour and alkylation with 27.5 M iodoacetamide in the dark for 20 minutes. The samples then underwent solid-phase extraction cleanup with an Oasis hydrophilic-lipophilic-balanced plate (Waters, Milford, MA), and the resulting samples were injected into an Orbitrap Fusion Lumos mass spectrometer (Thermo, Waltham, MA) coupled to an Ultimate 3000 RSLC-Nano liquid chromatography (LC) system (Thermo Fisher). Samples were injected onto a 75 mm internal diameter (ID) by a 75 cm long EasySpray column (Thermo Fisher) and eluted with a gradient from 0 to 28% buffer B over 90 minutes. Buffer A contained 2% (vol/vol) acetonitrile (ACN) and 0.1% formic acid in water, and buffer B contained 80% (vol/vol) ACN, 10% (vol/vol) trifluoroethanol, and 0.1% formic acid in water. The mass spectrometer was operated in positive-ion mode with a source voltage at 1.5 kV and an ion transfer tube temperature of 275°C. Mass spectrometry (MS) scans were acquired at a resolution of 120,000 in the Orbitrap instrument, and up to 10 tandem MS (MS/MS) spectra were obtained in the ion trap for each full spectrum acquired using higher-energy collisional dissociation (HCD) for ions with charges 2 to 7. Dynamic exclusion was set for 25 seconds after an ion was selected for fragmentation. Raw MS data files were analyzed using Proteome Discoverer v2.4 SP1 (Thermo Fisher), with peptide identification performed using Sequest HT, searching against the human protein databases from UniProt. Fragment and precursor tolerances of 10 ppm and 0.6 Da were specified, and three missed cleavages were allowed. Carbamidomethylation of Cys was set as a fixed modification, with oxidation of Met being set as a variable modification. The false discovery rate (FDR) cutoff was 1% for all peptides.

### Functional enrichment and pathway analyses

Normalized protein abundance was achieved by taking the sum of the raw abundance values per condition. Each value was multiplied by a scaling factor to equal the same total sum of protein abundances across samples. Bar graphs were generated using FunRich software v.3.1.3. We identified differentially abundant proteins in the chronic HOS group using a two-step process. First, a protein was deemed to be present in the control or treated group only if all three biological replicates of a group had non-zero abundance. Second, a threshold of >= 2 or <= 0.5 fold-change of normalized abundance was set to categorize enriched or depleted proteins, respectively. Pathway analysis was performed using g:Profiler to map differential candidates to the KEGG and CORUM databases and identify their functional impact^25^. Significantly enriched or depleted KEGG pathways were identified with an adjusted p-value of 0.05. For protein complexes, CORUM entries were filtered for those mapping to at least two differential proteins and a minimum 5% of all subunits to highlight multi-protein complexes enriched in the HOS sEVs. All statistical analyses and data visualization were performed using base packages and the tidyverse suite in the R statistical environment.

### Statistics

Statistical significance of densitometry quantitation on immunoblots, and concentrations of sEVs released from hTCEpi cells were tested by using an unpaired Student two-tailed t-test. A *P* value of <0.05 is considered statistically significant.

## Results

### Hyperosmolar stress increases sEV release and alters the proteomic cargo

One of the hallmark phenotypes of dry eye disease (DED) is hyperosmolarity of the precorneal tear film^26^. This increase in osmolarity results from either increased tear evaporation and/or reduced aqueous tear production. While the osmolarity of the human tear film is around 308 mOsm, this value is obtained from tears that are pooled at the inferior tear meniscus and may not reflect the actual osmolarity of tears spread across the corneal surface^27^. To determine the impact of HOS on sEV release, we developed a chronic subtoxic stress model to mimic the early stages of DED before the onset of cornea epithelial cell degeneration. We treated hTCEpi cells with 450 mOsm NaCl. Our lab has previously shown that this concentration is optimal for these studies^4^. As shown in Supplementary Figure S1A, hTCEpi cells exposed to HOS did not exhibit any abnormal morphological changes or signs of cell death compared to the isosmolar controls. To ensure that HOS induced oxidative stress, we performed immunoblotting analysis for Superoxide Dismutase (SOD1), an antioxidant (Suppl. Figure S1B). Densitometry confirmed a reduction in SOD1 levels in HOS hTCEpi cells (Suppl. Figure S1C), indicating an overall increase in cell stress.

Next, sEVs were isolated from normal and HOS hTCEpi cells by cushioned iodixanol density gradient ultracentrifugation. Immunoblotting confirmed the presence of several canonical sEV markers, including Tumor Susceptibility Gene 101 (TSG101), ALIX, CD9, and Annexin 2 (ANXA2) in the sEVs (Figure 1A and Suppl. Figure S3A). The absence of Calreticulin (CALR), an ER marker, confirmed the lack of cellular contamination in the sEV preparations. As shown in Figure 1 (A-E), there was a significant increase in specific sEV markers in the HOS condition, suggesting an increase in sEV release. We found a similar increase in the number of sEVs released from HOS hTCEpi cells (Figure 1F) using nanoparticle tracking analysis. There was no difference in size between sEVs released from isosmolar control and HOS hTCEpi cells (Figure 1G).

**Figure 1:**
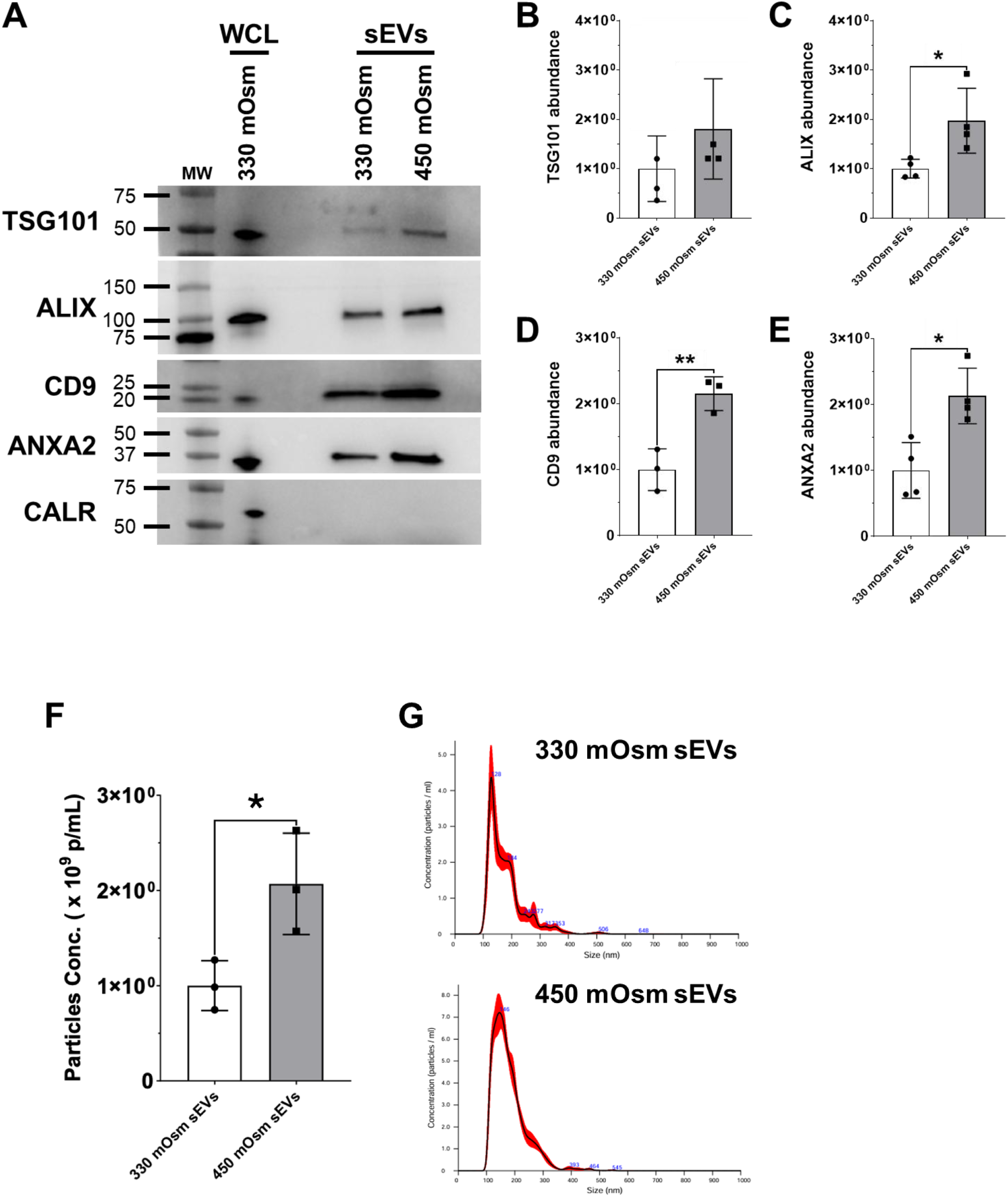
Hyperosmolar stress triggers an increase in sEV release. sEVs were isolated from isosmolar (330 mOsm) and HOS (450 mOsm) hTCEpi cells. **(A)** Immunoblotting confirmed the presence of established sEV markers TSG101, ALIX, CD9, and ANXA2. Calreticulin (CALR) was used as a control to confirm the absence of contamination in sEV samples. WCL: whole cell lysate (positive control), N=3. **(B-E)** Quantification of chemiluminescent immunodetection signal density on immunoblots, *p<0.05, ** p<0.01, t-test, N=3 for CD9, N=4 for all other immunoblots. **(F)** sEVs were isolated from isosmolar (330 mOsm) and HOS (450 mOsm) hTCEpi cells. NTA was used to quantify sEVs. We detected an increase in the number of sEVs released from HOS hTCEpi cells compared to control cells, *p<0.05, Unpaired t-test, N=3. **(G)** The size distribution of sEVs released from HOS hTCEpi cells and controls. Graphs show that sEVs are within the sEV range 30-150 nm in diameter^20^.

### Pathway analysis reveals a distinct proteomic signature in sEVs released from hyperosmolar-stressed corneal epithelial cells

Since sEVs release was increased in response to HOS, we next performed untargeted proteomics to examine cargo composition. Equal amounts of sEV protein were loaded and confirmed by silver staining (Suppl. Figure S2). A total of 822 proteins were detected, of which 778 proteins were shared by sEVs released from control and HOS hTCEpi cells (Figure 2A). In sEVs released from HOS cells, we identified 38 unique proteins. Consistent with our immunoblotting, we found an increase in the abundance of sEV markers TSG101, Annexin 5, Annexin 2, and Syntenin. (Table 1). There were only 6 unique proteins in sEVs released from control cells. We then performed Principal Component Analysis (PCA) on normalized protein abundance to assess variation between samples. We detected a distinct difference between sEVs released from HOS cells and control hTCEpi cells (Figure 2, B-C). To identify the biological processes affected by HOS, we analyzed the sEV proteome using KEGG pathway analysis (Figure 3). Interestingly, we found that HOS sEVs were enriched with proteins involved in endocytosis, pyruvate metabolism, cysteine, and methionine metabolism pathways (Figure 3A). Furthermore, to investigate whether functional protein complexes were packaged in sEVs, we mapped enriched and depleted proteins to the comprehensive resource of mammalian protein complexes (CORUM) database. We detected elevated levels of several subunits of the proteosome complex in HOS sEV cargo (Figure 3B). We then examined the proteosome complexes in detail and found that the Proteasome 20S Subunit Alpha (PSMA) 1, PSMA3, PSMA4, and PSMA6 were highly abundant in the cargo of sEVs released from HOS cells (Figure 4, A-D). It has previously been reported that proteasomes are packaged into sEVs when the cell is stressed and in certain disease states^28–31^. In contrast to these changes, focal adhesion and lysosome complexes were found to be depleted in the proteomes of HOS-sEVs (Figure 3A and Suppl. Figures S3 and S4). We also detected a lower abundance of desmosomal proteins in sEVs from HOS hTCEpi cells (Figure 5). We next examined several endocytosis and metabolism-associated markers (Figure 6). We found that Ras-related protein Rab-5C (RAB5C), Ras-related protein Rab-7A (RAB7A), and vacuolar protein sorting-associated protein 26A (VPS26A) were higher in abundance in sEVs derived from cells exposed to HOS (Figure 6, A-C). Metabolism-associated proteins such as Lactate Dehydrogenase A (LDHA), LDHB, Malate Dehydrogenase 1 (MDH1), and MDH2 were also found to be highly abundant in sEVs released from HOS cells (Figure 6, D-G).

**Figure 2:**
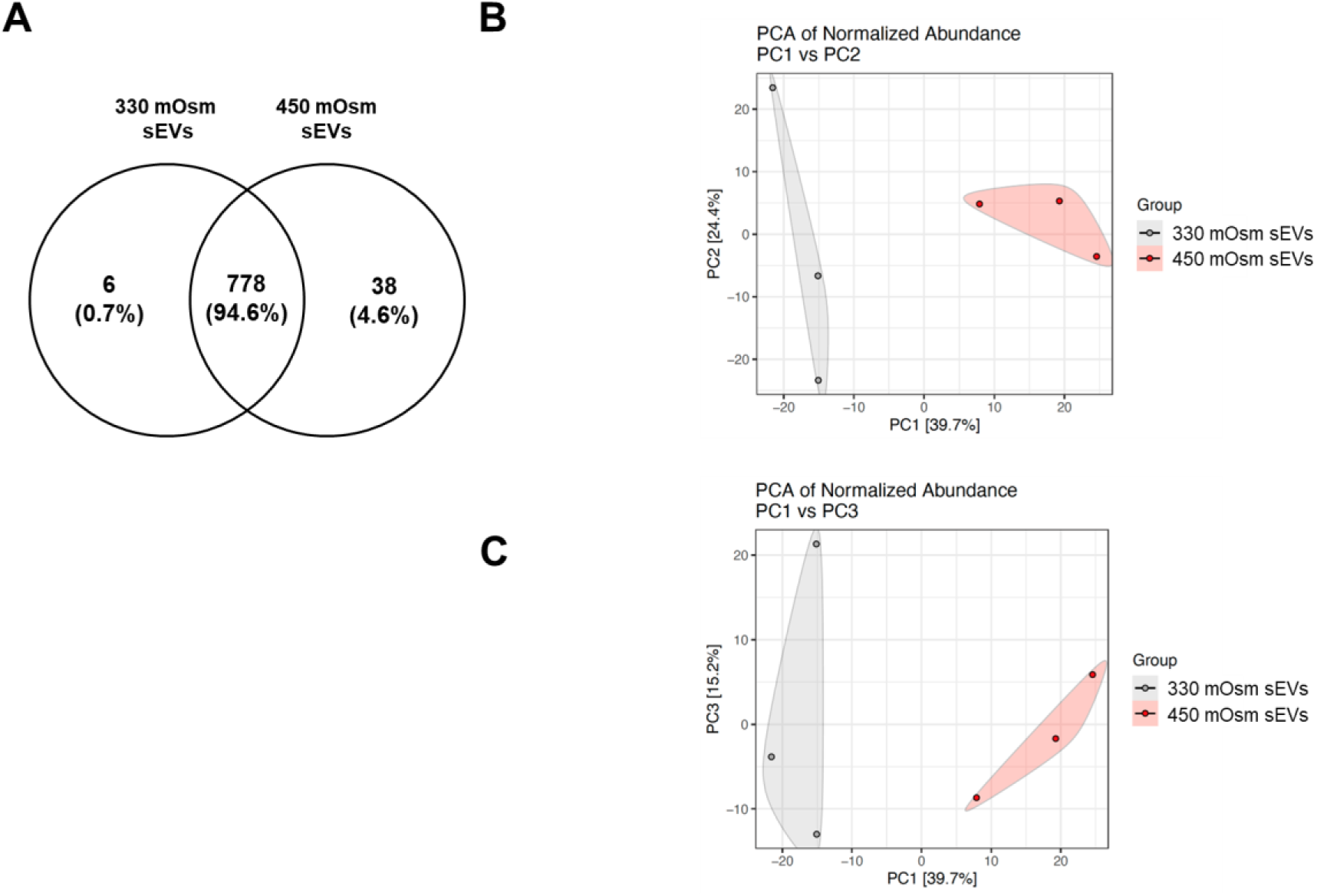
Protein profiling of sEVs released from normal and HOS hTCEpi cells reveals distinct proteome cargo based on HOS. **(A)** Venn Diagram showing shared and unique proteins detected in sEVs isolated from hTCEpi cells in isosmolar media (330 mOsm) and in hyperosmolar media (450 mOsm NaCl). **(B)** Principal Component Analysis (PCA) of normalized protein abundance showing PC1 vs PC2 blot. Each point represents an individual sample. N=3. **(C)** Principal Component Analysis (PCA) of normalized protein abundance showing PC1 vs PC3 blot. Each point represents an individual sample. N=3.

**Figure 3:**
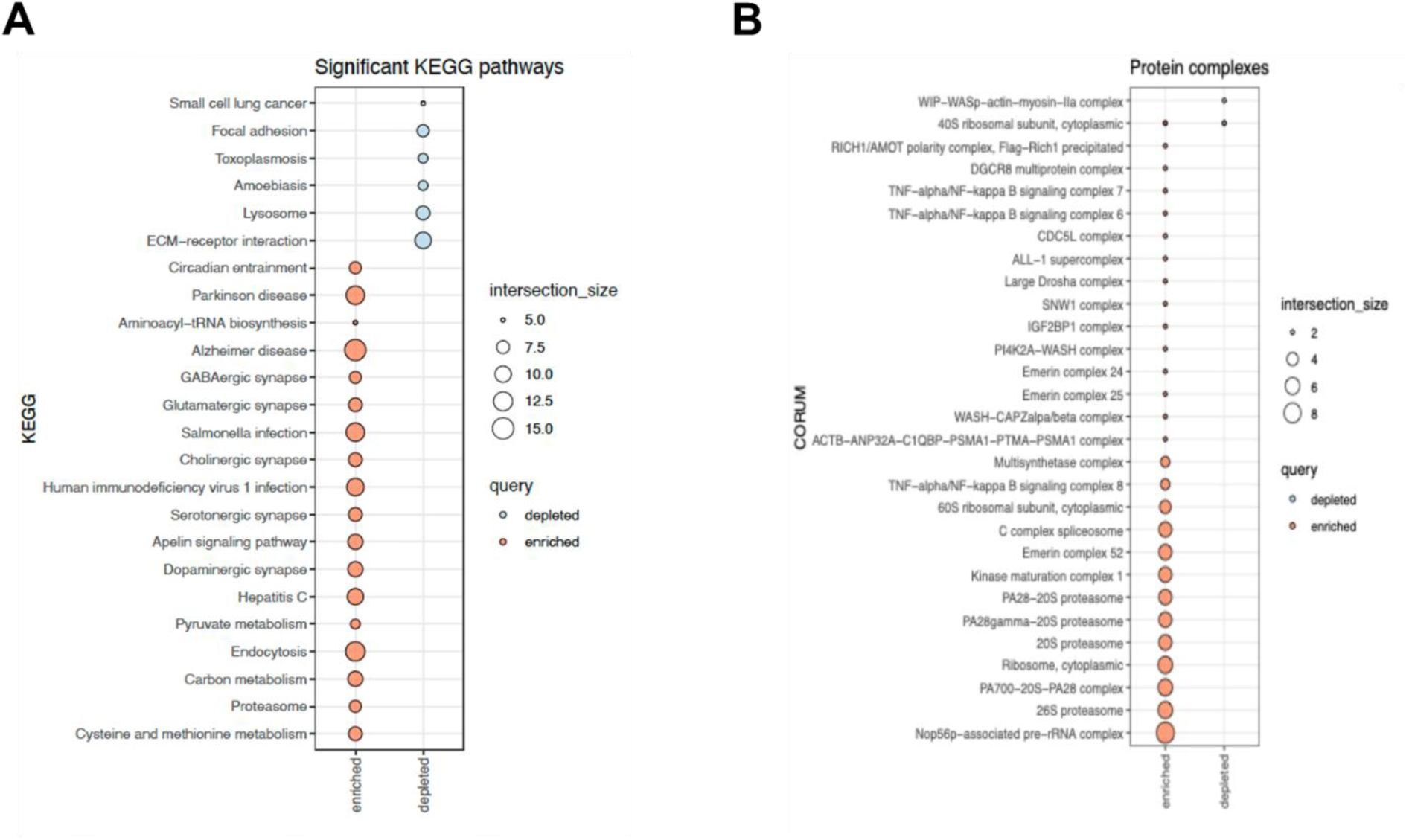
Biological pathway and protein complex enrichment analysis of sEVs released from isosmolar and hyperosmolar hTCEpi cells. **(A)** KEGG pathways enriched (red) and depleted (blue) in sEVs released from HOS hTCEpi cells. **(B)** Protein complexes enriched (red) and depleted (blue) in sEVs released from HOS hTCEpi cells.

**Figure 4:**
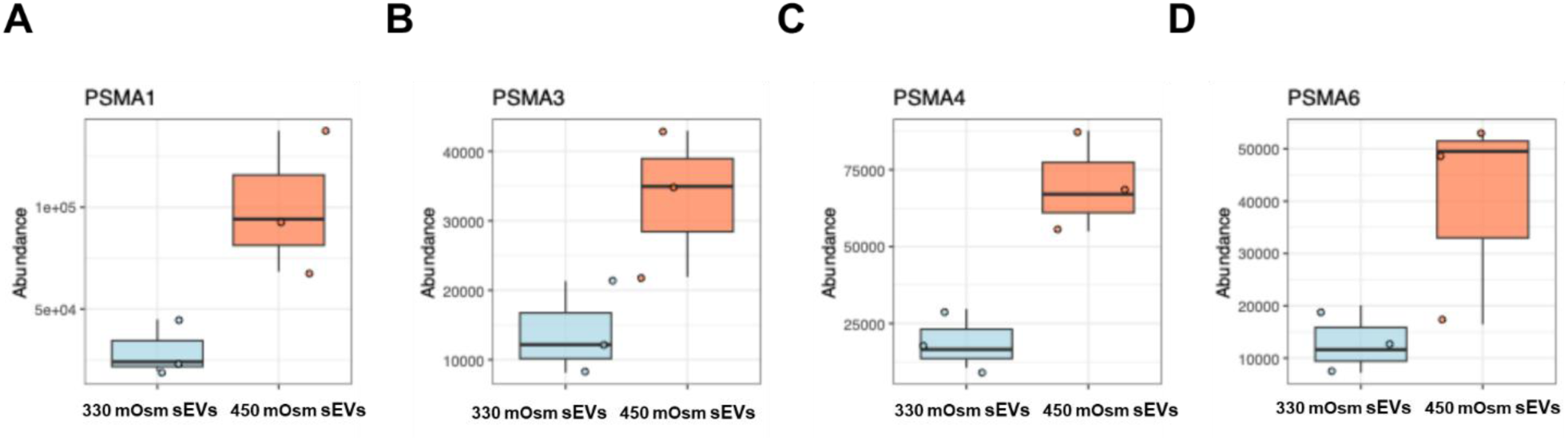
Specific protein complexes are enriched in sEVs released from HOS hTCEpi cells. **(A-D)** The top and bottom whisker of each boxplot represent where 25% of the data is located, and the horizontal line is the median. Protein complexes enriched (red) in sEVs from HOS cells compared to sEVs from isosmolar cells (blue).

**Figure 5:**
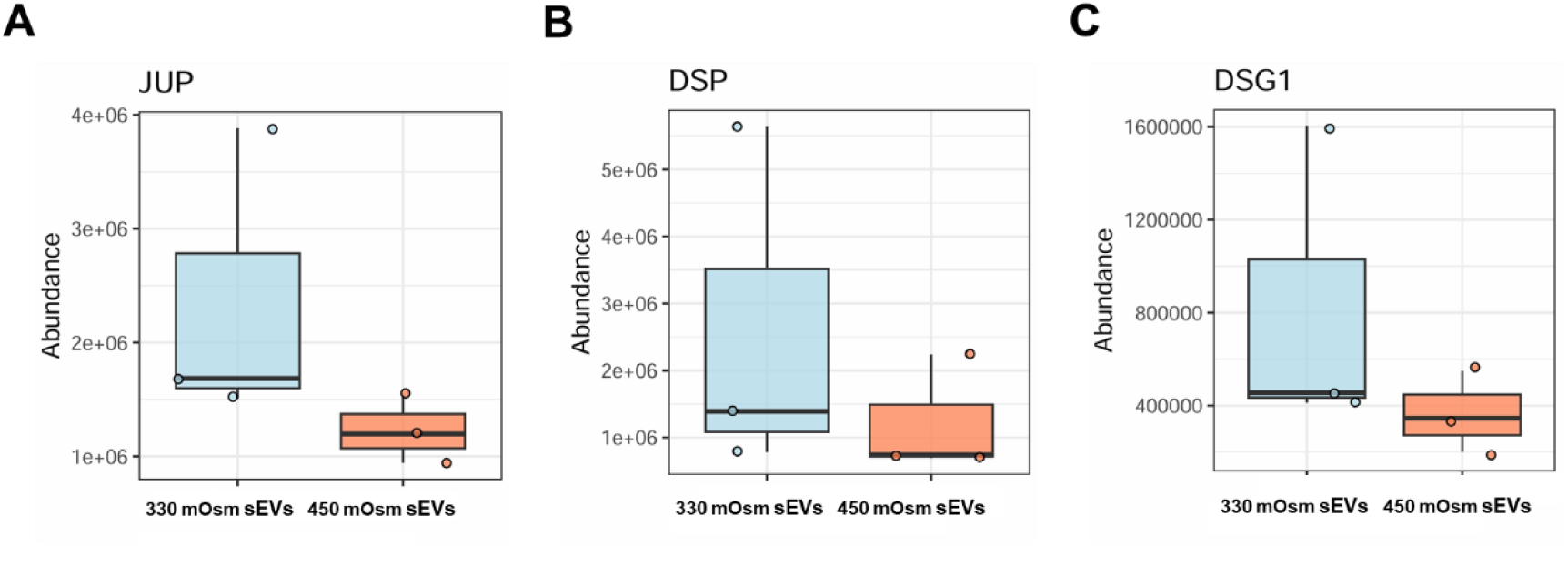
Elevated desmosome-associated proteins found in sEVs released from isosmolar control hTCEpi cells. The top and bottom whisker of each boxplot represent where 25% of the data is located, and the horizontal line is the median. **(A-C)** Desmosome proteins: JUP, DSP, and DSG1 were found in higher abundance in sEVs released from isosmolar control hTCEpi cells (blue) compared to sEVs released from HOS hTCEpi cells (red).

**Figure 6:**
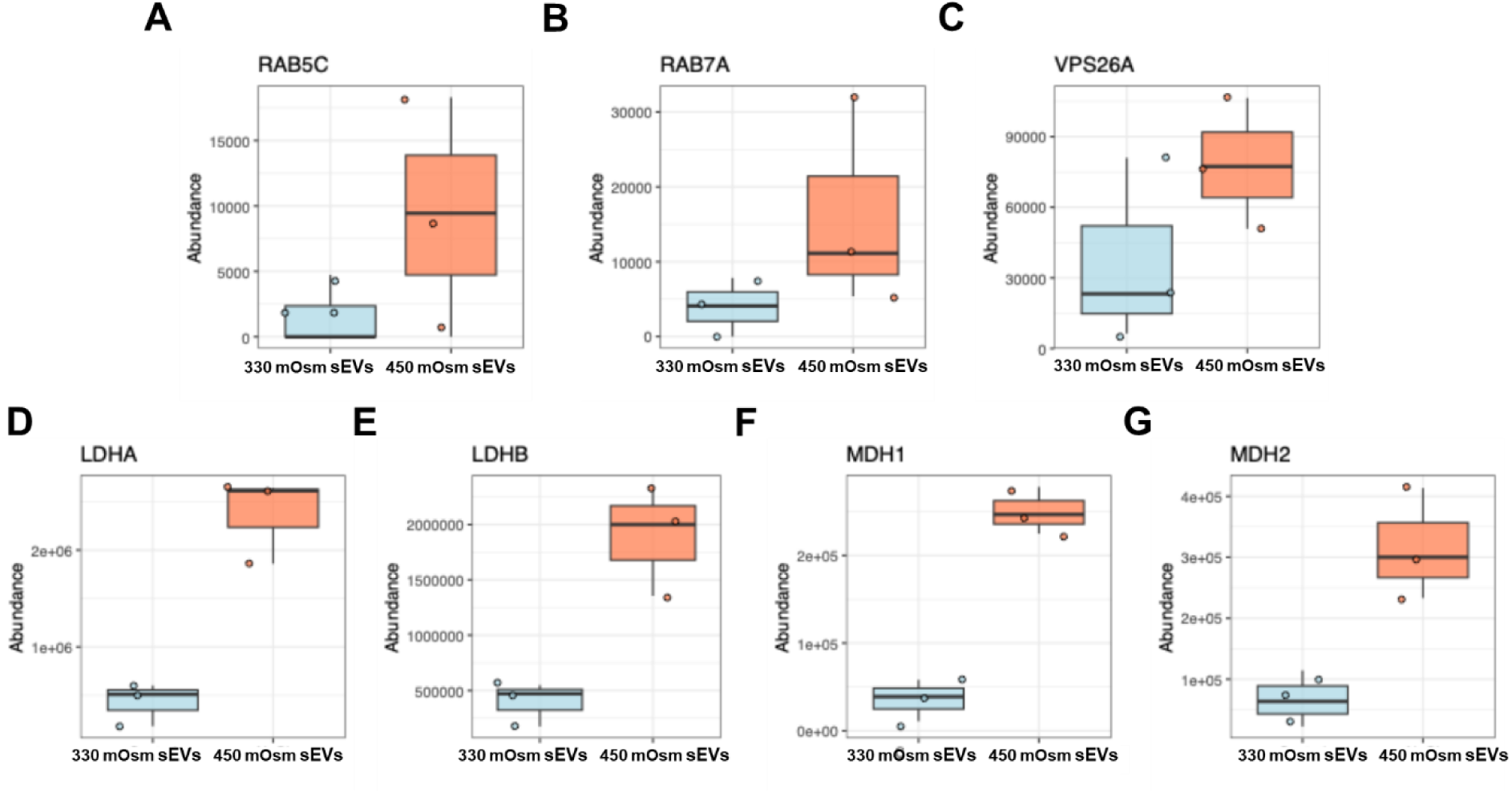
Detection of increased abundance levels of endocytosis and metabolism-associated proteins found in sEVs released from HOS hTCEpi cells. The top and bottom whisker of each boxplot represent where 25% of the data is located, and the horizontal line is the median. **(A-C)** Endocytosis proteins: RAB5C, RAB7A, and VPS26A were found in higher abundance in sEVs released from HOS hTCEpi cells (red) compared to sEVs released from isosmolar control hTCEpi cells (blue). **(D-G)** Metabolism-associated proteins such as LDHA, LDHB, MDH1, and MDH2 were found in higher abundance in sEVs released from HOS hTCEpi cells (red) compared to sEVs released from isosmolar control hTCEpi cells (blue).

**Table 1.** Metabolism-associated genes were increased in sEVs in response to hyperosmolar stress. The table below shows proteins in sEVs released from HOS hTCEpi cells compared to sEVs released from control cells with a fold change >2, N=3. The ranking of proteins is sorted from highest to lowest fold change.

| Protein | Gene | 330 mOsm sEV<br>Normalized<br>Abundance | 450 mOsm sEV<br>Normalized<br>Abundance | 450 mOsm/330 mOsm Fold<br>Change |
| --- | --- | --- | --- | --- |
| Phosphoglycerate mutase 1 | PGAM1 | 1.35 E04 | 1.00 E05 | 7.43 |
| Malate dehydrogenase, cytoplasmic | MDH1 | 3.60 E04 | 2.50 E05 | 6.95 |
| Ras-related protein Rab-5C | RAB5C | 1.57 E03 | 9.25 E03 | 5.87 |
| L-lactate dehydrogenase A | LDHA | 4.33 E05 | 2.37 E06 | 5.48 |
| Annexin A5 | ANXA5 | 2.47 E04 | 1.26 E05 | 5.08 |
| L-lactate dehydrogenase B | LDHB | 3.99 E05 | 1.90 E06 | 4.76 |
| Malate dehydrogenase,<br>mitochondrial | MDH2 | 6.68 E04 | 3.16 E05 | 4.72 |
| Ras-related protein Rab-7a | RAB7A | 3.97 E03 | 1.61 E04 | 4.05 |
| Apolipoprotein E | APOE | 2.48 E04 | 9.51 E04 | 3.84 |
| Syntenin | SDCBP | 1.41 E05 | 5.32 E05 | 3.77 |
| Annexin A2 | ANXA2 | 4.33 E06 | 1.35 E07 | 3.12 |
| Claudin-1 | CLDN1 | 8.32 E04 | 2.19 E05 | 2.63 |
| Alcohol dehydrogenase class 3 | ADH5 | 1.03 E04 | 2.65 E04 | 2.56 |
| Stomatin | STOM | 9.89 E05 | 2.46 E06 | 2.49 |
| Caveolae-associated protein 1 | CAVIN1 | 4.27 E04 | 9.09 E04 | 2.13 |
| Triosephosphate isomerase | TPI1 | 1.63 E05 | 3.22 E05 | 1.97 |
| Phosphoglycerate kinase 1 | PGK1 | 5.23 E05 | 9.29 E05 | 1.75 |

### Hyperosmolar stress triggers the release of sEVs that contain metabolic proteins

To confirm the increase in metabolic proteins, we validated the proteomic data by immunoblotting. We confirmed a significant increase in three proteins associated with cell metabolism. The first protein is Lactate Dehydrogenase A (LDHA, Figure 7, A-B), which plays a central role in the conversion of pyruvate to lactate and the oxidation of NADH to NAD+^32, 33^. We also found an increase in Phosphoglycerate Kinase 1 (PGK1, Figure 7, A-C). PGK1 is a key metabolic enzyme in glycolysis that generates ATP by converting 1,3-bisphosphoglycerate to 3-phosphoglycerate^34^. Finally, we found an increase in Malate Dehydrogenase 2 (MDH2, Figure 7, A-D). MDH2 is an enzyme involved in catalyzing the final tricarboxylic (TCA) cycle reaction, which converts malate to oxaloacetate, facilitating the continuity of the cycle and ATP production^35^. To determine whether the HOS-mediated increase in LDHA, PGK1, and MDH2 was associated with intracellular levels of these enzymes, we performed immunoblotting of whole cell lysates. While we detected a significant decrease in MDH2 levels in HOS hTCEpi cells (Figure 8, A&B), levels of PGK1 (Figure 8, A&C) and LDHA (Figure 8, A&D) were unchanged. Prior work in our lab has shown that HOS leads to an overall decrease in corneal epithelial cell metabolism^4^. Thus, the increase in metabolic proteins in HOS-sEVs suggests that HOS dismantles key components essential for major cell metabolic pathways and releases some of these components to the extracellular environment through the secretion of sEVs. These data further suggest that the presence of metabolic proteins in sEVs may represent early indicators of corneal epithelial cell dysfunction.

**Figure 7:**
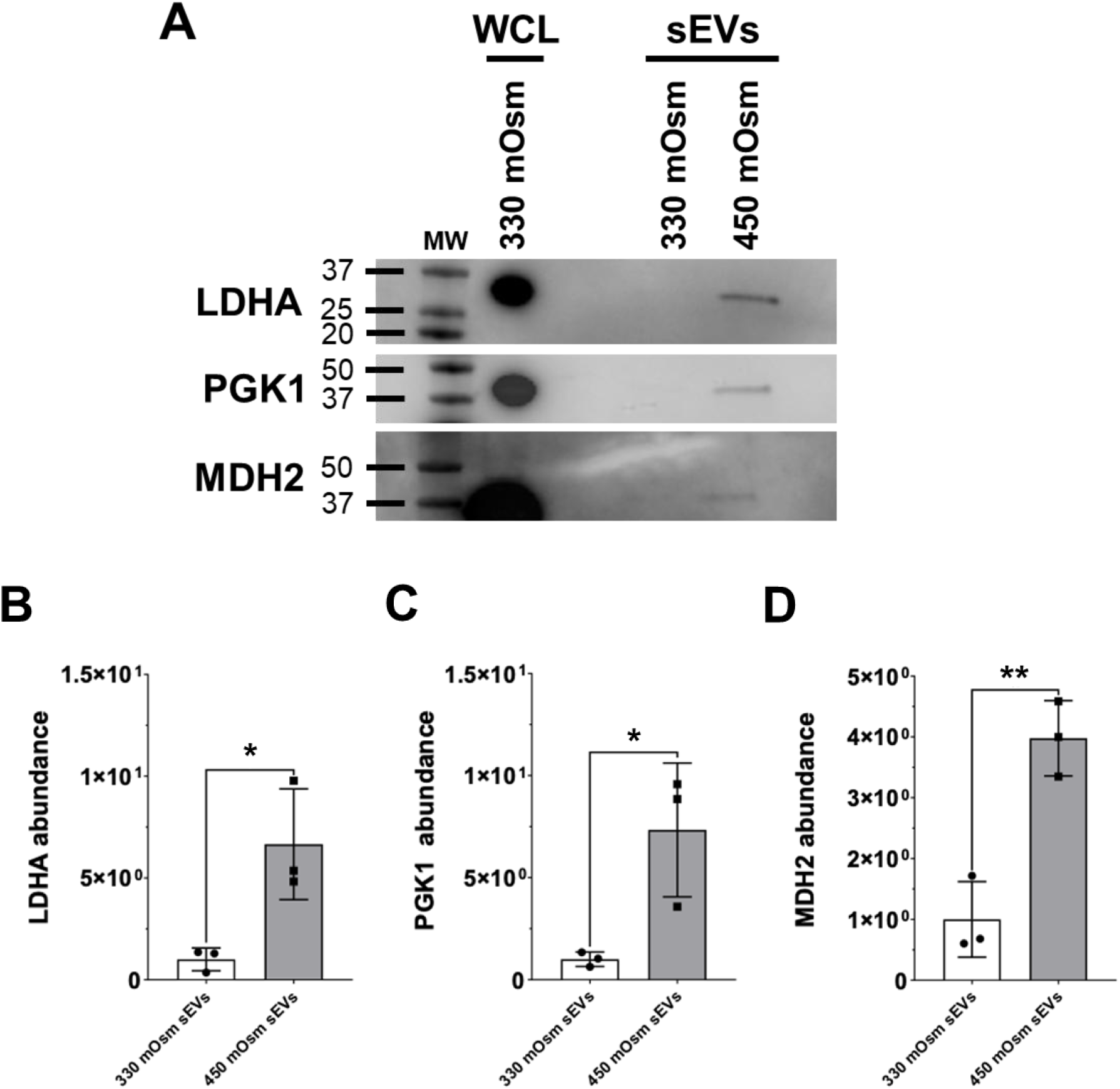
There is an increase in proteins associated with cell metabolism in sEVs released from HOS hTCEpi cells. To validate our proteomic data, we immunoblotted for LDHA, PGK1, and MDH2. **(A)** There was a significant increase in the abundance of LDHA, PGK1, and MDH2 in the cargo of sEVs released from HOS hTCEpi cells compared to isosmolar control cells. **(B-D)** Quantification of chemiluminescent immunodetection signal density on immunoblots, *p<0.05, **p>0.01, t-test, N=3.

**Figure 8:**
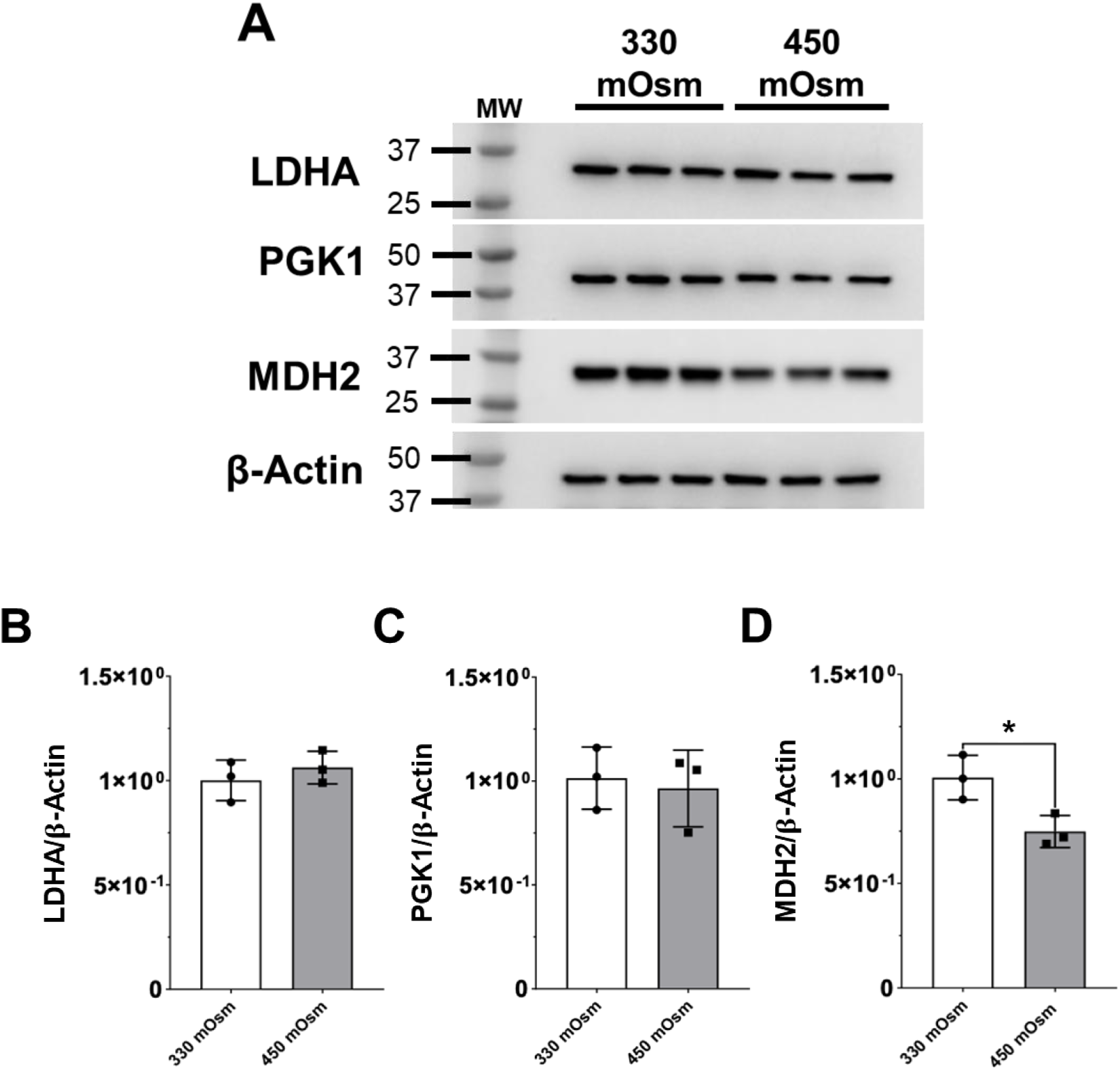
There is a decrease in expression of MDH2 in corneal epithelial cells exposed to HOS. To analyze the abundance of proteins associated with metabolism in control (330 mOsm) and HOS (450 mOsm) hTCEpi cells, we immunoblotted for LDHA, PGK1, and MDH2. **(A)** Immunoblot analysis of isosmolar control and HOS whole-cell lysates showed no difference in LDHA and PGK1 protein levels. MDH2 expression was reduced in HOS hTCEpi cells. **(B-D)** Quantification of chemiluminescent immunodetection signal density on immunoblots, *p<0.05, t-test, N=3. β-actin was used as a loading control.

### Hyperosmolar stress perturbs intercellular junctions in corneal epithelial cells

Desmosomes, gap junctions, and adherens junctions are the major structural components of the corneal epithelium. While desmosomes impart mechanical strength, gap junctions and adherens junctions play a key role in cell-to-cell communication and signaling. While the tight epithelial barrier is disrupted in DED, there are limited studies on the effect of HOS on corneal epithelial integrity. Interestingly, analysis of our proteomic data (Table 2 and Figure 5, A-C) revealed a higher abundance of desmosomal proteins in the cargo of sEVs released from isosmolar control cells. To validate these findings, we performed immunoblotting for three desmosomal protein markers, Desmoplakin 1 (DSP1), Junctional Plakglobin (JUP), and Desmoglein 1 (DSG1). However, unlike our mass spectrometry findings, we were unable to detect a change in the level of these junctional markers in sEV cargo released from HOS and control cells (Figure 9, A-D). Thus, immunoblotting lacked the sensitivity that was needed to detect these small differences in protein abundance. We next examined the expression of these proteins in whole cell lysates. Unlike sEVs, we found that JUP was significantly decreased in lysates from HOS cells (Figure 10, A&B). There were no detectable changes in DSP or DSG1 in cell lysates between normal and control cells (Figure 10, A and C-D). These findings suggest that in response to HOS, there is a retention of junctional proteins in an attempt to maintain tissue integrity, despite the attenuation of cell metabolism. The decrease in intracellular expression of JUP may further represent an early sign of junctional compromise.

**Figure 9:**
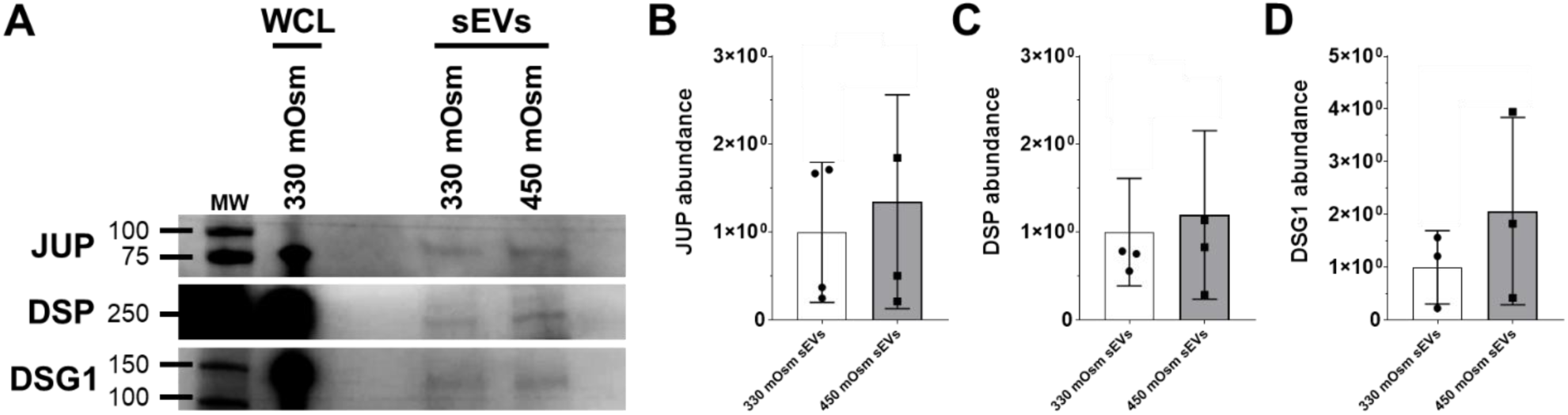
There was no significant change in desmosomal protein abundance in the cargo of sEVs released from hTCEpi cells exposed to hyperosmolar stress. To analyze the abundance of proteins associated with desmosomes, we used JUP, DSP, and DSG1. **(A)** We detected the presence of desmosomal proteins in the cargo of sEVs released from hTCEpi cells under isosmolar and HOS conditions. **(B-D)** Quantification of chemiluminescent immunodetection signal density on immunoblots, *p<0.05, t-test, N=4, except DSG1 N=3.

**Figure 10:**
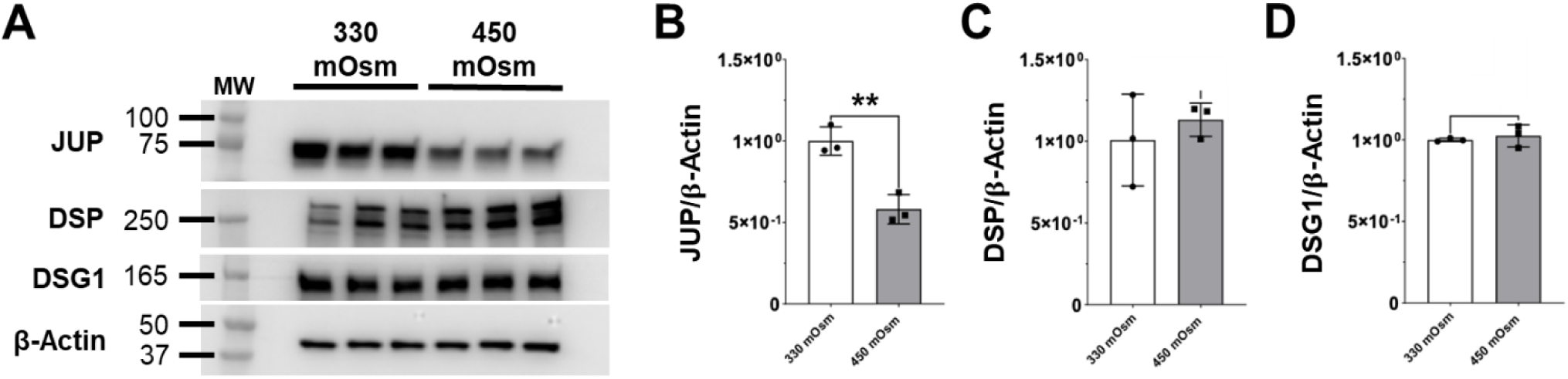
Expression of desmosomal proteins DSP and DSG1 are unchanged during exposure to hyperosmolar stress. To analyze the abundance of proteins associated with adhesion in control (330 mOsm) and HOS (450 mOsm) hTCEpi cells, we immunoblotted for JUP, DSG1, and DSP. **(A)** Western blot analysis of control and HOS hTCEpi whole-cell lysates showed no difference in DSG1 and DSP protein levels. However, we detected a decrease in JUP levels in HOS CECs. **(B-D)** Quantification of chemiluminescent immunodetection signal density on immunoblots, **p<0.01, t-test, N=3. β-actin was used as a loading control.

**Table 2.** Desmosome-associated proteins are increased in sEVs released from control corneal epithelial cells. The table below shows proteins present in sEVs from HOS hTCEpi cells compared to sEVs released from control cells with a fold change >2, N=3. The ranking of proteins is sorted from highest to lowest fold change.

| Protein | Gene | 330 mOsm sEV Normalized Abundance | 450 mOsm sEV Normalized Abundance | 330 mOsm/450 mOsm Fold Change |
| --- | --- | --- | --- | --- |
| Galectin-3 binding protein | LGALS3BP | 9.30 E05 | 2.29 E05 | 4.06 |
| Galectin-7 | LGALS7 | 8.24 E04 | 2.10 E04 | 3.92 |
| Complement C3 | C3 | 8.97 E05 | 2.88 E05 | 3.11 |
| Dermcidin | DCD | 1.52 E06 | 5.45 E05 | 2.78 |
| Prolactin-inducible protein | PIP | 1.07 E05 | 3.93 E04 | 2.72 |
| Dystroglycan 1 | DAG1 | 7.92 E04 | 2.99 E04 | 2.64 |
| Progranulin | GRN | 2.01 E05 | 7.62 E04 | 2.64 |
| Desmocollin-1 | DSC1 | 3.72 E05 | 1.47 E05 | 2.53 |
| EGF-containing fibulin-like extracellular matrix protein | EFEMP1 | 6.47 E05 | 2.68 E05 | 2.42 |
| Desmoglein-1 | DSG1 | 8.24 E05 | 3.65 E05 | 2.26 |
| Desmoplakin | DSP | 2.60 E06 | 1.22 E06 | 2.13 |
| Peroxidasin | PXDN | 1.64 E05 | 7.90 E04 | 2.08 |
| Calmodulin-like protein 5 | CALML5 | 2.54 E05 | 1.23 E05 | 2.07 |
| Lysosome-associated membrane glycoprotein 1 | LAMP1 | 3.60 E05 | 1.79 E05 | 2.02 |
| Olfactomedin 4 | OLFM4 | 1.47 E05 | 7.44 E04 | 1.98 |
| Junction plakoglobin | JUP | 2.36 E06 | 1.23 E06 | 1.92 |

## Discussion

This study investigated the release and cargo composition of sEVs when corneal epithelial cells are exposed to a HOS. Importantly, we found an increase in EVs released from HOS cells. The size distribution, measured using NTA, was consistent with the reported size of sEVs. In addition, immunoblotting also indicated an increase in sEV release, as increases in known sEV markers, ALIX, CD9, and ANXA2, were detected. To confirm that our HOS cells were exposed to subtoxic chronic stress, we also performed immunoblotting for SOD1, an antioxidant that is commonly decreased in conditions of chronic stress. As expected, expression levels of SOD1 were decreased in our model.

The most notable finding in this study is the enrichment of proteins in sEVs that are involved in cell metabolism. This increase in the release of metabolic proteins coincides with our prior data describing the HOS-mediated impairment in corneal epithelial cell metabolism.^36, 37^ In addition, we also found an increase in proteins associated with the proteasome. The function of the proteosome is intricately tied to metabolism, since the proteosome requires ATP to enable proper function. Moreover, a reduction in cellular proteasomal activity has been associated with several different types of disease^28–31^. Together, these findings indicate that HOS triggers an increase in proteotoxic stress. While autophagy is increased in HOS in attempts to recycle damaged proteins, these data suggest that the increase in misfolded proteins may overwhelm the cell’s lysosomal degradative and proteasome pathways, leading to an increase in sEV release^4^.

Maintenance of cell metabolic activity is crucial for cellular homeostasis^38^. Alterations of cell metabolic pathways are suggested to be the driving force behind disease states such as cancer, cerebral aneurysms, and heart failure^39–41^. In our HOS model, we detected an increase in the release of LDHA, PGK1, and MDH2 via sEVs, which are likely excess proteins being secreted due to decreased metabolic activity in HOS cells. In Alzheimer’s Disease, it has been previously shown that MDH2 levels are increased following the induction of oxidative stress^42^. Indeed, our SOD1 levels confirmed that the cells were subject to oxidative stress in our model. PGK1 has been identified as a potential biomarker for breast cancer, where it has been shown to regulate glycolytic activities^43^. Of high relevance to the current study, increased levels of MDH1 and MDH2 have been identified in tears from DED patients compared to normal healthy controls^44^. That same study further suggested that tear levels of MDH2 may provide a new way to monitor response to therapy in DED^44^. In contrast to metabolic proteins, desmosomal proteins were not increased in HOS sEVs. Instead, except for JUP, they were retained intracellularly. This suggests that the cell may be working to maintain some level of junctional integrity. Future studies are needed to determine the significance of this observation.

EVs have been previously identified in human tears, and current studies point towards the use of tear EVs as potential biomarkers for certain diseases.^37, 45, 46^ EVs have also been examined for their potential role in corneal wound healing.^13, 14, 47^ *In vitro* human cornea epithelial cell culture models have shown that sEVs released from corneal epithelial cells, stromal fibroblasts, and endothelial cells accelerate wound closure^13^. Other studies have explored the effects of corneal epithelial cell-derived sEVs on corneal wound healing using *in vivo* rat models^14^. EVs derived from mesenchymal stem cells have also been shown to exert beneficial effects on corneal wound repair and ocular damage in dry eye disease, further opening the door for possible therapeutic applications.^48, 49^ This has led to a surge in studies evaluating whether custom-loaded EVs can be designed to enhance the treatment of various ocular diseases.^16, 50^ We have also reported on distinct changes in the proteomic and metabolic composition of EVs released by bacteria and host cells during infection by *Pseudomonas aeruginosa.* Importantly, we have found that the EVs released during infection are important modulators of inflammation and bacterial invasion^23, 24, 51^. The ability of sEVs to mediate inflammation in infectious and non-infectious corneal disease is a burgeoning field and requires significantly more work.

A key strength of the current study is the method of sEV isolation. To isolate sEVs that were free of cellular contaminants, we used a cushioned iodixanol buoyant density gradient ultracentrifugation (C-DGUC). The majority of studies analyzing the proteome and transcriptome in EVs released by corneal cells have been achieved by using ExoQuick (PEG precipitation) and differential ultracentrifugation (DUC)^13, 14, 52, 53^. Both ExoQuick and DUC methods have been shown to capture a heterogeneous population of large and small EVs, lipoproteins, and various amounts of soluble protein contaminants^54^. In contrast to this, previously published EV studies have shown that the C-DGUC method, as used here, provides a pure yield of sEVs ^52, 55, 56^.

In conclusion, this is the first study to demonstrate an increase in the release of cell metabolic and proteasomal proteins via corneal epithelial cell-derived sEVs in response to HOS. Moreover, the HOS sEV protein signature that we have described, before the onset of overt cell morphological dysfunction, suggests that sEVs may serve as early indicators of dry eye disease. Studies are now underway to determine whether these metabolic indicators are present in sEVs isolated from human tear fluid and how their composition in sEVs may be used to reflect disease severity in clinical practice.

