## Supplemental information for "Hyperosmolar stress promotes the release of small extracellular vesicles containing metabolic proteins from corneal epithelial cells"

### Supplementary Figures and Tables

**Supplementary Table S1. Unique proteins in sEVs released from HOS-treated hTCEpi cells.** This table shows proteins that were only present in the sEVs released from HOS-treated hTCEpi cells and expressed zero peptides in sEVs released from isosmolar control cells.

| Protein | Gene |
|---|---|
| Annexin A3 | ANXA3 |
| 14-3- protein eta | YWHAH |
| Alpha-soluble NSF attachment protein | NAPA |
| Solute carrier family 26 member 6 | SLC26A6 |
| Glyoxylate reductase/hydroxypyruvate reductase | GRHPR |
| Ras-related protein Rab-35 | RAB35 |
| ATP synthase subunit alpha, mitochondrial | ATP5F1A |
| Serine/threonine-protein phosphatase 2A catalytic subunit beta isoform | PPP2CB |
| Delta(3,5)-Delta (2,4)-dienoyl-CoA isomerase, mitochondrial | ECH1 |
| CYFIP-related Rac1 interactor B | CYRIB |
| ADP-ribosylation factor 3 | ARF3 |
| Proteasome subunit alpha type-2 | PSMA2 |
| Aldo-keto reductase family 1 member A1 | AKR1A1 |
| 26S proteasome regulatory subunit 10B | PSMC6 |
| Matrin-3 | MATR3 |
| BTB/POZ domain-containing protein KCTD12 | KCTD12 |
| Proteasome activator complex subunit 1 | PSME1 |
| CCN family member 2 | CCN2 |
| Syntaxin-4 | STX4 |
| Nucleolar and coiled-body phosphoprotein 1 | NOLC1 |
| Acetyl-CoA acetyltransferase, mitochondrial | ACAT1 |
| PDZ and LIM domain protein 1 | PDLIM1 |
| Ras-related protein Rap-2b | RAP2B |
| Casein kinase II subunit alpha | CSNK2A1 |
| Serine/arginine-rich splicing factor 1 | SRSF1 |
| Phosphatidylinositol transfer protein beta isoform | PITPNB |
| Insulin-like growth factor 2 mRNA-binding protein 2 | IGF2BP2 |
| Sulfide:quinone oxidoreductase, mitochondrial | SQOR |
| Eukaryotic translation initiation factor 3 subunit M | EIF3M |
| S-formylglutathione hydrolase | ESD |
| Peroxiredoxin-6 | PRDX6 |
| Lamin-B1 | LMNB1 |
| Developmentally-regulated GTP-binding protein 1 | DRG1 |
| DNA replication licensing factor MCM6 | MCM6 |
| Phosphatidylethanolamine-binding protein 1 | PEBP1 |
| Unconventional myosin-Ie | MYO1E |
| Palladin | PALLD |
| 26S proteasome regulatory subunit 8 | PSMC5 |

**Supplementary Table S2. Unique proteins in sEVs released from isosmolar control hTCEpi cells.** This table shows proteins that were only detected in sEVs released from isosmolar control hTCEpi cells and expressed zero peptides in sEVs released from HOS cells.

| Protein | Gene |
|---|---|
| Receptor-type tyrosine-protein phosphatase S | PTPRS |
| Epiplakin | EPPK1 |
| Immunoglobulin gamma-1 heavy chain | IGG1 |
| Immunoglobulin kappa constant | IGKC |
| Dolichyl-diphosphooligosaccharide--protein glycosyltransferase subunit 1 | RPN1 |
| Desmocollin-2 | DSC2 |

**Supplementary Figure S1: There was no loss in cell viability or changes in corneal epithelial cell morphology under chronic subtoxic HOS.** Microscopy images of hTCEpi cells at confluence in isosmolar (330 mOsm) and HOS (450 mOsm NaCl) media at day 0, day 1, and day 5. **A)** The characteristic polygonal cell shape was maintained under HOS conditions. **B)** Western blot confirmed for SOD1 confirmed that hTCEpi cells cultured in HOS were subject to oxidative stress. **C)** Quantification of chemiluminescent immunodetection signal density on immunoblots. SOD1 expression levels were normalized to β-actin, *$p<0.05$, t-test, N=3.

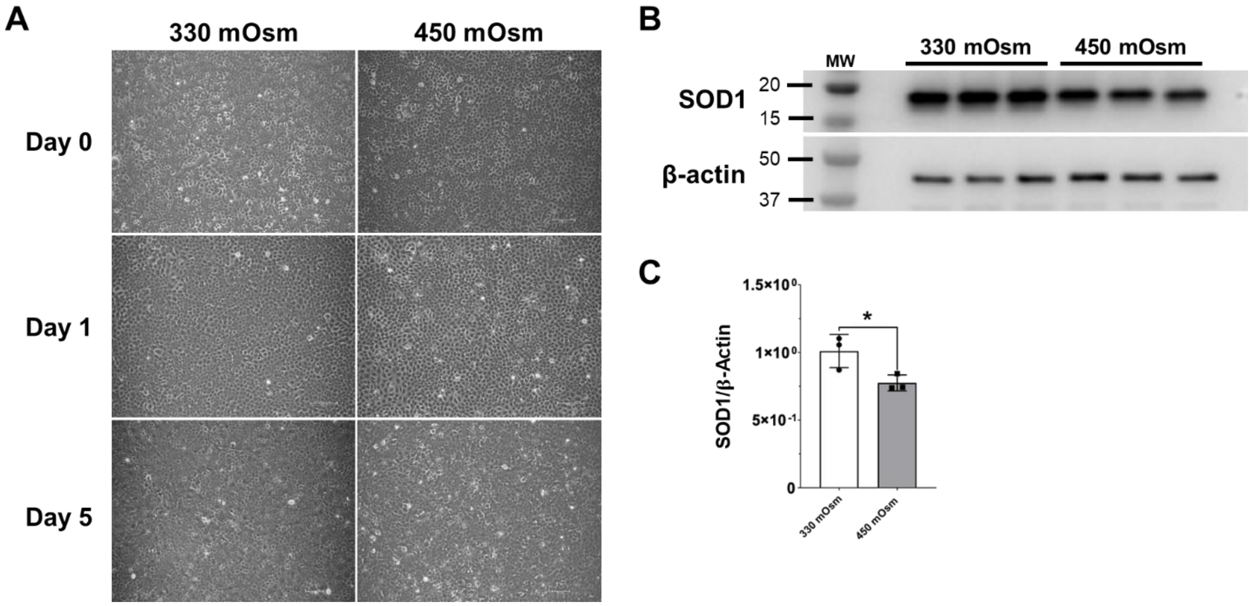

**Supplementary Figure S2: Equal loading of sEVs was confirmed by silver stain.** Silver stain image of sEVs isolated from hTCEpi cells cultured in isosmolar (330 mOsm) and HOS (450 mOsm NaCl) conditions. N=2.

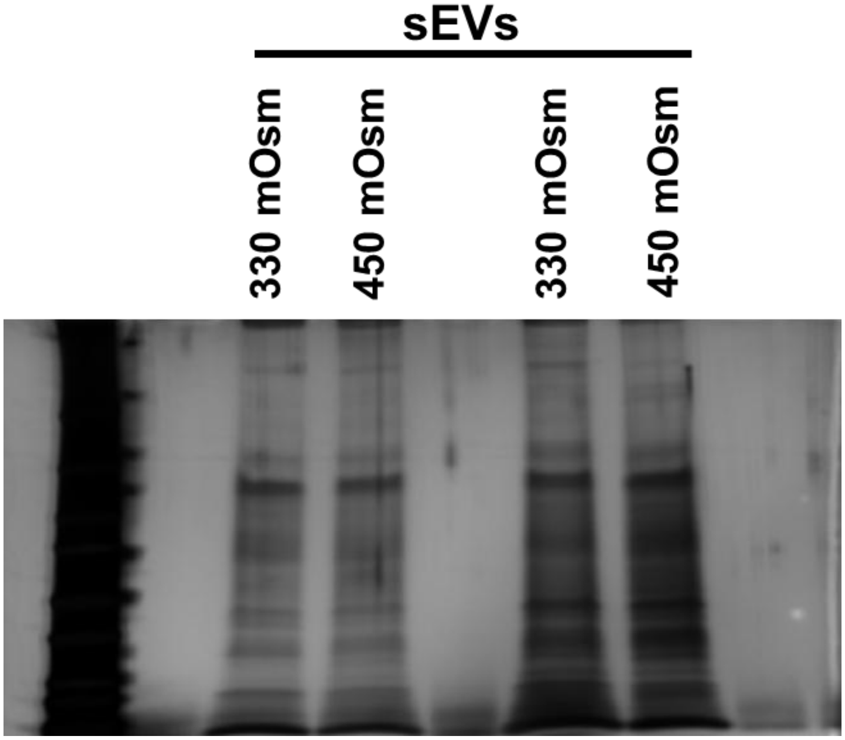

**Supplementary Figure S3: Protein cargo in sEVs released from HOS corneal epithelial cells indicates a disruption in cell metabolism.** We performed FunRich Analysis on proteomic data from three replicate sets of sEVs released from isosmolar control and HOS-hTCEpi cells. All proteins had a fold change >2 in the sEVs released from HOS hTCEpi cells compared to sEVs from control cells. **A)** We detected an increase in protein markers for sEVs in the HOS condition. **B)** We found an increase in proteins associated with cell and protein metabolism in the cargo of sEVs released from HOS cells. Proteins involved in cell growth and maintenance were higher in sEVs released from control cells.

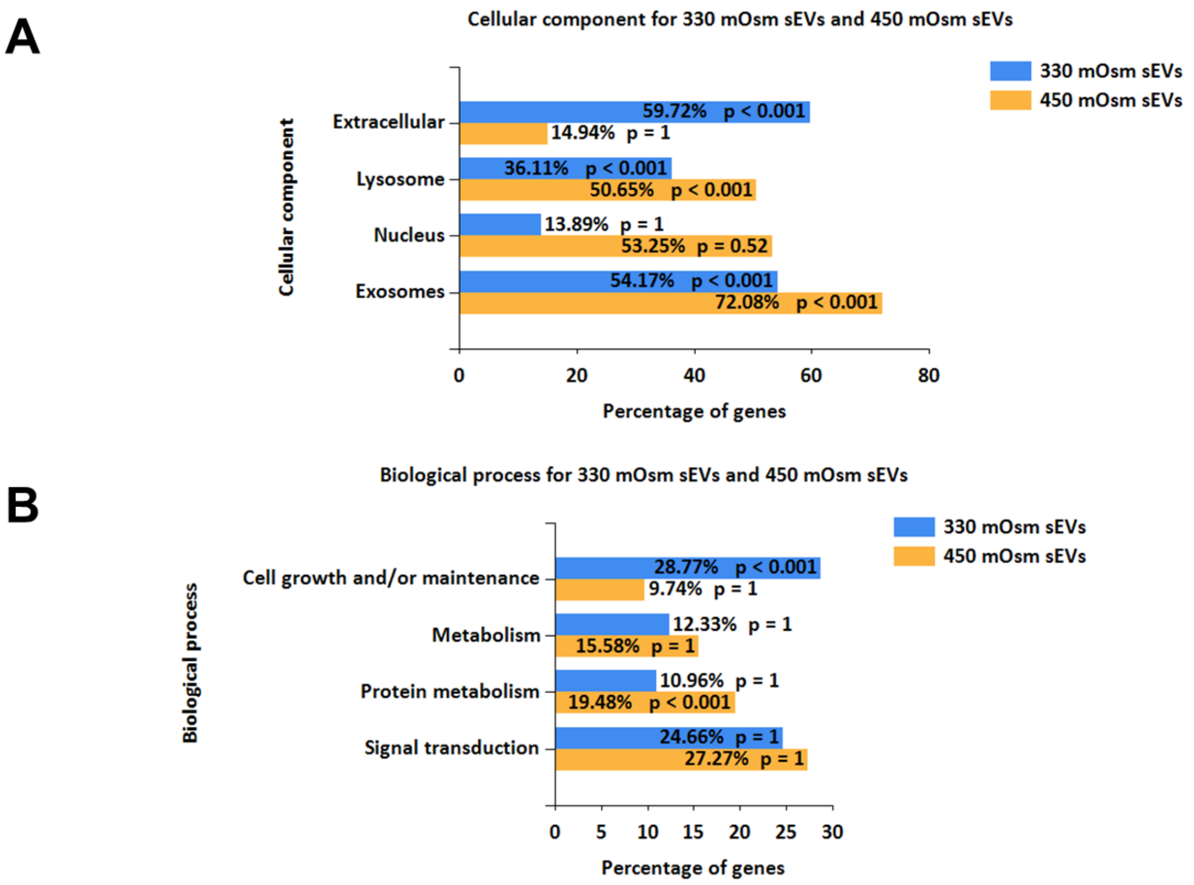

**Supplement Figure S4: There was a decrease in the abundance of lysosomal proteins in sEVs released from HOS-treated hTCEpi cells.** The top and bottom whisker of each boxplot represent where 25% of the data is located, and the horizontal line is the median. **A-D)** Lysosomal proteins: LAMP1, LAMP2, SCARB2, and GAA were decreased in abundance in sEVs released from HOS hTCEpi Cells (orange) compared to sEVs released from isosmolar control cells (blue).

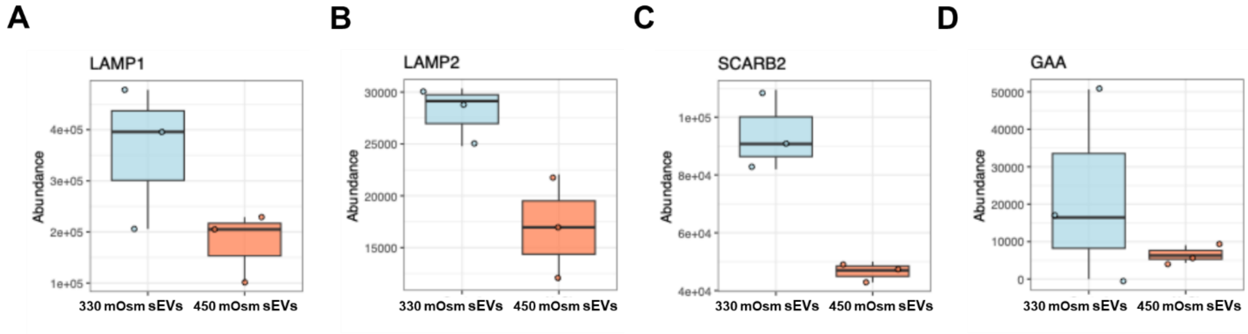
